# The association between transcriptional regulation of macrophage differentiation and activation and genetic susceptibility to inflammatory bowel disease

**DOI:** 10.1101/2022.12.14.520373

**Authors:** Claire L O’Brien, Kim M Summers, Natalia M Martin, Dylan Carter-Cusack, Rasel Barua, Ojas VA Dixit, David A Hume, Paul Pavli

## Abstract

The abundant macrophage population of the intestinal lamina propria turns over rapidly and is replaced by blood monocytes. The differentiation and survival of resident intestinal macrophages depends upon signals from the macrophage colony-stimulating factor receptor (CSF1R). The response of human monocyte-derived macrophages (MDM) grown in macrophage colony-stimulating factor (CSF1) to bacterial lipopolysaccharide (LPS) has been proposed as a model for the differentiation and adaptation of monocytes entering the intestinal lamina propria. We hypothesized that dysregulation of this response leads to susceptibility to chronic inflammatory bowel disease (IBD). To address this hypothesis we analyzed transcriptomic variation in MDM from affected and unaffected sib pairs/trios from 22 IBD families and 6 healthy controls. There was no overall or inter-sib distinction between affected and unaffected individuals in basal gene expression or the stereotypical time course of the response to LPS. However, the basal or LPS-inducible expression of individual genes including inflammatory cytokines and many associated with IBD susceptibility in genome-wide association studies (GWAS) varied by as much as 100-fold between subjects. Extreme independent variation in the expression of pairs of HLA-associated transcripts (*HLA-B/C, HLA-A/F* and *HLA-DRB1/DRB5*) was associated with HLA genotype providing a novel explanation for the HLA association with disease susceptibility. The relationship between single nucleotide variant (SNV) genotype and gene expression at other loci was weaker and inconsistent suggesting that much of the variation arises from the integration of multiple *trans*-acting effects. For example, expression of *IL1B* at 2 hrs of LPS treatment was significantly associated with local SNV genotype and with peak expression of IL23A at 7 hrs. By contrast, there was no evidence of association between peak *IL6* mRNA at 7hrs, *IL6*-associated SNV genotype or *IL1B* at 2 hrs. Our results support the view that gene-specific dysregulation in macrophage adaptation to the intestinal milieu provides a plausible explanation for genetic susceptibility to IBD. The analysis also suggests that the molecular basis of susceptibility is unique to each individual which may contribute to variation in the precise environmental trigger, the consequent pathology and response to treatment.

**Author summary:** Cells of the innate immune system called macrophages are abundant in the wall of the gut, providing a first line of defense against potential pathogens. These cells must also avoid an inappropriate or excessive response to the abundant microbial population (the microbiome) of the intestine. We have previously proposed that genetic differences between individuals in macrophage adaptation to the unique environment of the intestine underlie genetic susceptibility to inflammatory bowel disease (IBD). In this study we developed a model of the adaptation of macrophages and used that model to identify surprisingly extreme variation in the response amongst a cohort of affected and unaffected siblings in families with IBD. The response did not distinguish affected individuals from their unaffected siblings. Our results support the view that each individual within IBD-susceptible families carries a unique set of genetic variants of large effect that together predispose to uncontrolled gut inflammation in the face of an environmental trigger.

## Introduction

The major chronic inflammatory bowel diseases (IBD), Crohn’s disease (CD) and ulcerative colitis, clearly have significant heritability. The strongest associations are with the human major histocompatibility gene complex (HLA), contributing up to 30% of shared risk amongst siblings for CD and even greater for ulcerative colitis [1, 2]. In terms of mechanism, there is a clear consensus that IBD arises from an aberrant interaction between cells of the immune system in the intestinal mucosa and the luminal microbiota [3]. Several rare monogenic early-onset forms of IBD have provided some mechanistic insights [4, 5]. Modern population genome-wide association studies (GWAS) have identified hundreds of additional non-HLA variants associated with significantly increased risk [3, 6–8]. Collectively, this genetic variation still explains only 20-30% of the heritability. The missing heritability has been attributed in part to limitations in sample size and other constraints on GWAS and to complex interactions between genes and environment. The genomic intervals identified by GWAS contain multiple genes and few SNVs are likely to be functional. To extend association analysis, whole genome sequencing has been applied to identify rare and/or causal variants within expression quantitative trait loci (eQTL) [9–11]. After sequencing 100 candidate genes in 6600 CD patients and 5500 controls, Momozawa *et al.* [10] concluded that > 10-fold greater sample sizes would be required to demonstrate causality. Accordingly, Sazonovs *et al.* [11] analyzed whole exome or whole genome sequences of >30,000 CD patients and 80,000 controls to implicate coding variation in an additional 10 genes in susceptibility.

The strength and weakness of GWAS is the lack of a hypothesis that can focus analysis on a smaller set of genomic loci to obviate the tyranny of appropriate correction for multiple testing. There have been numerous hypothesis-driven candidate gene analyses providing evidence of associations between disease and SNVs associated with inflammatory mediators. For example, Richard *et al.* [12] identified a disease-associated functional regulatory variant linked to reduced expression of *TNFSF15* (TL1A) which has been implicated in gut inflammation. Recently, Klunk *et al.* [13] proposed that the medieval plague epidemic led to selection for variants affecting immune response gene expression. They highlighted a SNV linked to expression of the *ERAP2* gene in macrophages. The same SNV was previously associated with IBD risk [14]

The intestinal lamina propria contains an abundant population of resident tissue macrophages which turn over constantly and are replaced by recruitment of blood monocytes. Following their entry to the mucosal niche, monocytes must rapidly modulate their response to the products of microorganisms to avoid inflammatory activation [15, 16]. The differentiation of lamina propria macrophages depends upon macrophage colony-stimulating factor (CSF1) and these cells are rapidly depleted by administration of anti-CSF1R antibody [17]. Accordingly, the differentiation of human monocytes *in vitro* in response to CSF1 and their subsequent response to bacterial agonists such as lipopolysaccharide (LPS) provides a model of differentiation of monocyte-derived macrophages in the ileum and colon [18].

Based upon extensive gene expression analysis in monocytes and macrophages produced with the FANTOM5 consortium, loci associated with IBD were strongly and specifically enriched for promoters that were regulated during monocyte differentiation or activation [18]. Furthermore, amongst known IBD susceptibility loci, the vast majority contained promoters that were regulated during the CSF1-dependent monocyte-macrophage transition and/or in response to LPS [18, 19]. These analyses also indicated that even amongst the most supported IBD susceptibility loci, genes in the same genomic region as the highlighted candidate are either co-regulated in monocytes (e.g. *SNX20* and *CYLD* with *NOD2*) or provide more plausible candidates (e.g. *INPP5D* within the *ATG16L1* locus) [18].

Siblings presumably share some components of both genotype and environment. It has been known for many years that the relative risk of developing CD is greater amongst the siblings of affected individuals. However, for monozygotic twins shared risk is estimated at only 15-30% and concordance even amongst dizygotic twins (who may share a more similar environment compared to siblings) may be <5% [20, 21]. In the current study we have performed gene expression analysis of monocyte-derived macrophages (MDM) and their response to LPS in a cohort of sib-pairs discordant for IBD disease. The extended cultivation of monocytes in CSF1 *in vitro* to produce MDM provides a common environment to reduce possible direct effects of disease status on monocyte function and enrich for detection of effects of genotype. Previous studies have identified allele-associated variation in regulated gene expression in monocytes and monocyte-derived macrophages amongst healthy individuals and associated with autoimmune disease susceptibility [22–26] Network analysis of the monocyte data [22] identified *trans*-acting impacts of variation in expression of *IFNB1* and downstream transcription factors IRF7 and IRF9 [27].

The genome of each affected individual with IBD must contain only a subset of the hundreds of disease-associated genetic variants that have been identified in population-based analyses and may also contain private variants of larger effect. If disease onset occurs purely in response to an environmental trigger that unaffected siblings have avoided (which is strongly implicated by the low level of concordance amongst siblings) then the entire cohort of affected individuals and first degree relatives will be strongly enriched for disease-associated variants regardless of disease status. Alternatively, unaffected siblings may be discordant for risk-associated variants and consequently at lower genetic risk of disease. In this study we demonstrate that MDM from both affected and unaffected individuals within families affected by IBD display extreme variation in expression of individual genes and classes of genes that may predispose to dysregulation of the response to microbial challenge and hence affect risk of developing IBD in response to an environmental trigger.

## Results

Monocyte-derived macrophages were prepared from a total of 56 individuals, including affected and unaffected sib pairs/trios from 22 IBD families and a set of 6 healthy individuals with no known family history of IBD, including one sib-pair. The response of human MDM to LPS is a sequential cascade of transient induction and repression of sets of transcripts [18]. Based upon the previous dense time course analysis using Cap analysis of gene expression (CAGE) tag sequencing [18], MDM populations were sampled at zero time, and 2, 7 and 21 hours after addition of a maximal effective concentration (100ng/ml) of *Salmonella Minnesota* Re595 LPS, a pure TLR4 agonist [28, 29]. The chosen time points represent the peaks of expression of early, mid and late response genes, also applied in previous comparative analysis of mouse and human responses [28] and multiple other mammalian species [29]. RNA-seq was performed on mRNA extracted at each time point and initially quantified using Kallisto as described previously [29]. The patient metadata is provided in **Table S1A** and the complete dataset organized by individual and family is provided in **Table S1B.**

To overview the data, we initially performed a network analysis using Biolayout [30]. A sample-to-sample network shown in **Figure 1A** (left panel) demonstrates that samples from each of the time points tend to group together, with a transition across the network from pre-LPS to 21 hours post-LPS reflecting the progressive temporal profile of the response. This is consistent with the global changes in gene expression elicited in response to LPS [18] but there was no class separation between affected and unaffected individuals (**Figure 1A** (right panel)). **Figure 1B** shows the main element of the gene-to-gene network graph based on the entire dataset, and **Table S2** shows the cluster gene lists alongside the average gene expression profiles of each cluster. Cluster 1, the largest, contains 1903 transcripts that were expressed constitutively and relatively invariant across the entire dataset with a small decline with time in response to LPS. This cluster comprises mainly house-keeping genes, but also includes macrophage-enriched transcription factors (*SPI1, CEBPA/B*) and surface markers (*ITGAM, ITGAX*). The consistent detection of these transcripts provides a data quality control and a reference point for much greater variation amongst other transcripts. A second large cluster, Cluster 4 (401 transcripts) is even less variable between individuals and highly-enriched for mitochondria-related transcripts and ribosomal protein genes. Cluster 12 (146 transcripts) was highly variable between individuals at 0 time and down-regulated by LPS. This cluster includes transcription factors *ETV5, MAFB*, *MEF2C, NFXL1*, *RB1* and multiple macrophage-enriched differentiation markers and cell surface receptors (*AIF1, CD163, CD28, CD302, CD4, CLEC10A, CSF1R, FOLR2, GAS6/7, GPR34, GSDMA, INPP5D, MERTK, MPEG1, MRC1, TLR7, TYROBP, STAB1, TNFRSF11A, VSIG4*). Their correlated expression indicates that there is significant variation in CSF1-induced differentiation amongst individuals. ETV5 [31], MAFB [32] and MEF2C [33] have each been identified as regulators of monocyte-macrophage differentiation, and variation in their expression (in each case >10-fold in the 0 time samples) likely contributes to the co-regulation of this cluster of genes. Many of these genes (including *MAFB* and *CSF1R*) form part of what was classified as a gut resident macrophage signature in single cell-RNA-seq analysis of IBD lesions [34]. There are several large clusters that reinforce previous analysis of the progressive temporal profile of the response of human MDM to LPS [18]. Transcripts in Cluster 11 (161 transcripts) peaked at 2 hours and then declined rapidly. They include genes encoding the key feed forward activators, *TNF, IFNB1* and *IFNL1*, transcriptional regulators (e.g. *ATF3, FOSL1, KLF6, KLF7, MYC, NFATC1, NFE2L2, NFIL3, NFKBIZ*, *NR4A3*) and many negative feedback regulators (e.g. *DUSP1, GPR183, NFKBIA, OTUD1, TNFAIP3* and *ZFP36L2*) that serve to constrain the response. Transcripts within Cluster 9 (209 transcripts) were also induced by 2 hours, but the expression declined less rapidly and was retained at 7 hours. This cluster includes additional transcription factors *IRF1, IRF8*, several interferon target genes (*IFIT2*), inducible cytokines (*IL1B, IL12A*) and chemokines (*CXCL1, CCL3, CCL4*).

**Figure 1.**
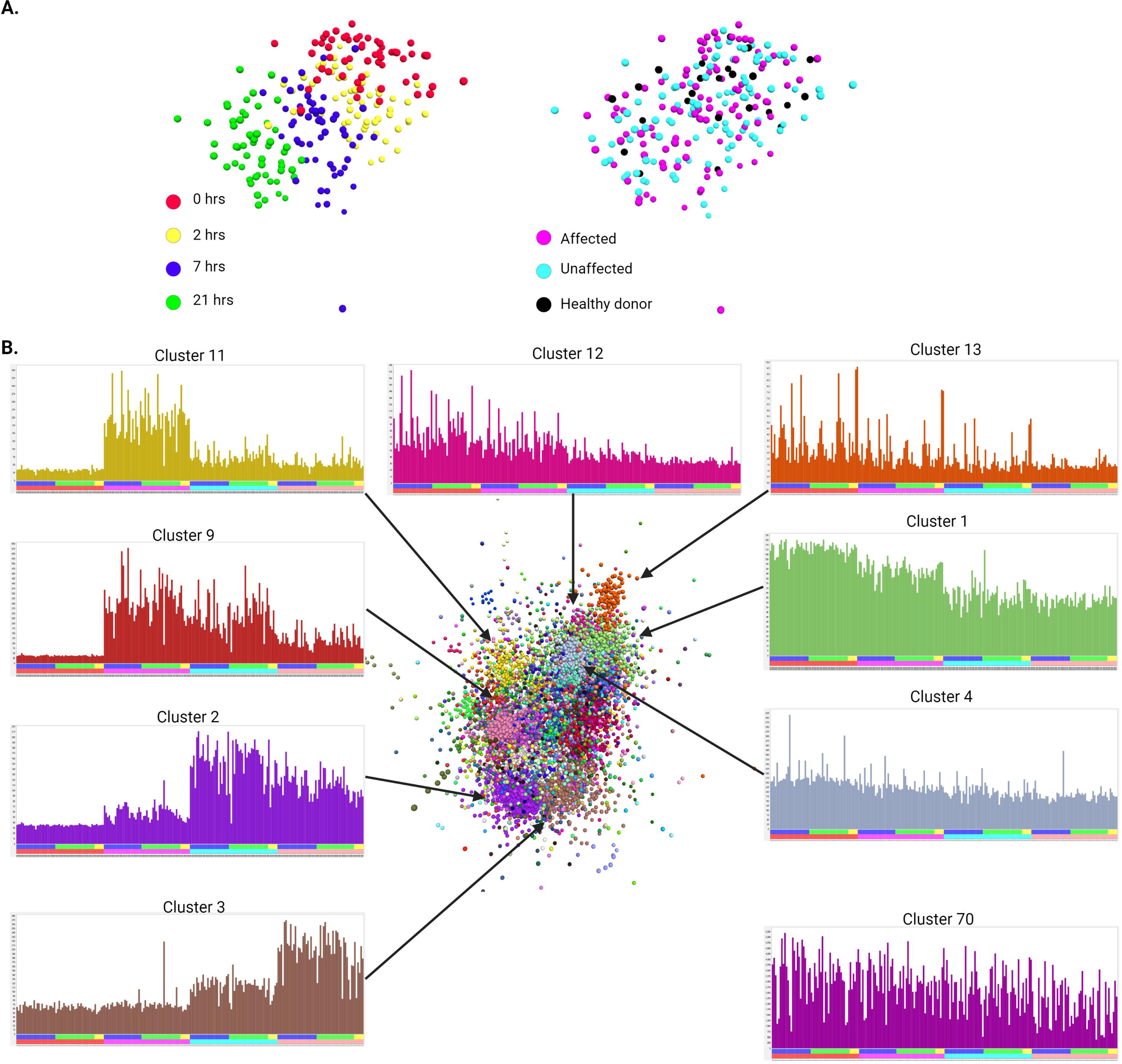
Network analysis of gene expression in macrophages stimulated with LPS. Gene expression profiles from RNA-seq analysis of the time course of the LPS response of MDM from 56 individuals (primary data in **Table S1**) were subjected to network analysis using Biolayout as described in Materials and Methods. Figure 1A*. Sample-to-sample network.* Each sphere represents a sample (individual time point). Edges between them (Pearson correlations of ≥ 0.87) have been omitted for clarity. Note the clear separation of the samples based upon time of incubation (left) but not on disease status (right). Figure 1B*. Gene-to-gene network.* Each sphere represents a gene. Edges between them (Pearson correlations of ≥ 0.7) gave been omitted for clarity and only the main element of the network is shown. Spheres are coloured by their membership of a cluster (MCL inflation value 2.2), Graphs show the averaged expression of genes in key clusters; colours of the columns match those in the network graph. Y axis – gene expression in CPM. X axis – the samples are ordered by time point and disease status; each column represents a single sample. Upper bar – disease status (blue: affected (Crohn’s disease or ulcerative colitis); green: unaffected sibling of an affected individual; yellow: healthy donor with no family history of inflammatory bowel disease). Lower bar – time point (red: 0 hrs; pink: 2 hrs; turquoise: 7 hrs; salmon: 21 hrs). The profiles for all clusters are provide in **Table S2.** Arrows indicate the position of the cluster in the network. The position of Cluster 70 is not indicated as it forms a small separate element not linked to the largest element shown here.

The large Cluster 2 (819 transcripts) groups the set of transcripts that was maximally induced at 7 hours post-stimulation and declined by 21 hours. It includes inducible transcription factors (*BATF, BATF2, BATF3, IRF9, NFE2L3, STAT2,3,4*) and further interferon target genes (e.g. *IFIT1, ISG15, MNDA, MX1, OAS1*). Similarly, Cluster 3 (445 transcripts) contains transcripts that were inducible by 7 hours but continued to increase, peaking at 21 hours in the response. This cluster also contains transcripts encoding transcriptional regulators (*CEBPB, IRF6, RELB, STAT1*) and a distinct cohort of chemokines, surface receptors (e.g *CLEC4E*) and interferon-inducible transcripts including the receptor *IFNAR2* and *IFI6, IFI27, IFITM1/2/3.* It is notable that *IRF7,* which also formed part of an IFN-inducible regulon in monocytes responding to LPS [27, 35] was not found in any cluster. Indeed, by contrast to the induction reported in monocytes *IRF7* was induced by LPS in MDM derived from only 9/56 individuals. On the other hand, *IRF1* (which may be affected by an autoimmune disease risk variant in monocytes [24]) was strongly induced by LPS to a similar extent in MDM in all individuals.

Although blood monocytes are predominantly post-mitotic, CSF1 is able to promote proliferation in a subset of cells and this response is inhibited by LPS [36]. Accordingly, Cluster 13 (104 transcripts) contains the key cell cycle transcription factors (*E2F1/E2F2/E2F8, FOXM1, MYBL2*) and numerous structural and regulatory genes that are induced specifically in S phase and mitosis [37]. The expression of transcripts in this cluster was relatively low, and down-regulated by LPS, but clustering arises because of the substantial variation between individuals at the earlier time points.

Cluster 70 (17 transcripts) contains all of the mitochondrial encoded transcripts, clustered because of the extensive and correlated variation between individuals. This observation is consistent with evidence for high levels of variation in expression of these transcripts in human populations [38]. No corresponding variation was detected in nuclear-encoded mitochondrial genes.

### Variation in the temporal profile of expression of key cytokine genes

Many pro-inflammatory cytokines produced by macrophages are targets for successful therapeutic intervention in IBD and/or are affected by regulatory variants that are in turn associated with susceptibility to inflammatory disease. Our focus in this study was based on the idea that affected individuals and their siblings provide a discovery cohort to identify extreme regulatory variation. As noted above, the majority of transcripts varied by less than 2-fold amongst the 50 individuals from IBD families at each of the 4 time points. To identify the most highly variable transcripts, in **Tables S1C-F** the genes have been ranked in order of maximum/minimum each time point. At each of the three time points post LPS treatment (2hrs, 7hrs, 21hrs) there were around 100 transcripts that varied more than 100-fold in expression between individuals (**Table S1D-F**). These include each of the cytokines mentioned above, and numerous chemokine genes.

In some cases, the variation reflects temporal differences. **Figure 2** shows the time courses of regulation of selected cytokines (*IL1A, IL1B, IL10, IL12B, IL23A, IL6, TNF* and *TNFSF15*) across the whole dataset. The transient induction of *TNF* with a peak at 2hrs and of *IL12B* at 7hrs was relatively consistent between individuals apart from a small number of outliers with low or sustained expression (**Figure 2, Table S1B**). For other transcripts, the pattern of regulation varied. For example, in 9 individuals, *IL6* mRNA peaked at a relatively low level at 2 hrs then declined, where in the majority, peak expression at a much higher level occurred at 7 hrs. In the majority (35/56), *IL1B* peaked at 2 hrs then declined rapidly, but in 21/56 *IL1B* mRNA was sustained or increased further at 7 and 21 hrs.

**Figure 2.**
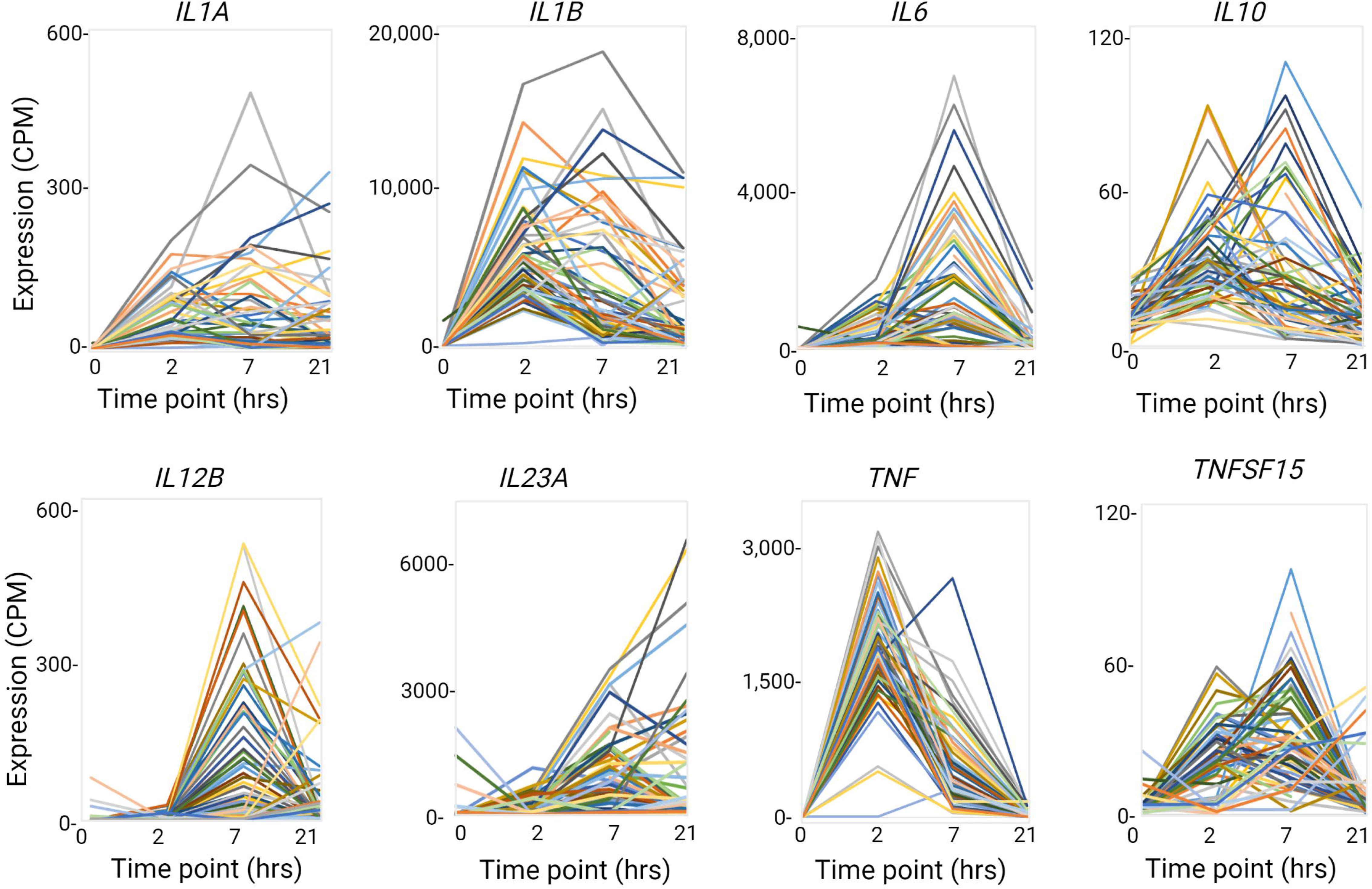
Individual variation in patterns of transcriptional regulation of key cytokine genes in LPS-stimulated MDM. Gene expression profiles for the indicated cytokine genes were extracted from the RNA-seq analysis of the time course of the LPS response of MDM from 56 individuals (primary data in **Table S1B-F**). Each line joins the 4 time points from a single individual.

IL23, a dimeric protein comprised of IL12B and IL23A, is a therapeutic target in IBD [39, 40]. **Figure 2** shows that *IL23A* and *IL12B* were each strongly induced by LPS but with distinct temporal profiles. In the cluster analysis, *IL23A* forms part of the small hypervariable **Cluster 118** (10 transcripts) whereas *IL12B* was not correlated with any other transcript at the threshold of r = 0.7. This observation raises the possibility that some individuals are IL23-deficient through a failure to coordinate the expression of the two subunits. **Figure 3A** shows that the expression of the genes encoding the two subunits is significantly correlated across the whole dataset. However, there are some individuals where one gene is very highly expressed while the other is low (**Figure 3B**). In addition, the peak expression of *IL23A* is an order of magnitude higher than expression of *IL12B.* IL23A may also form dimers with EBI3 to produce an alternative effector of the IL23R, although there is some doubt as to whether this occurs in humans [41]. Unlike *IL12B* and *IL23A*, *EBI3* forms part of the late response **Cluster 3** and was amongst the most highly-induced transcripts with relatively little variation amongst individuals. EBI3 is also a dimer partner with IL27A to form IL27. *IL27* mRNA was also LPS inducible and varied greatly between individuals, but unlike the dimer partner, it is part of **Cluster 2** (peaking at 7hrs) and is a known interferon target gene [42].

**Figure 3.**
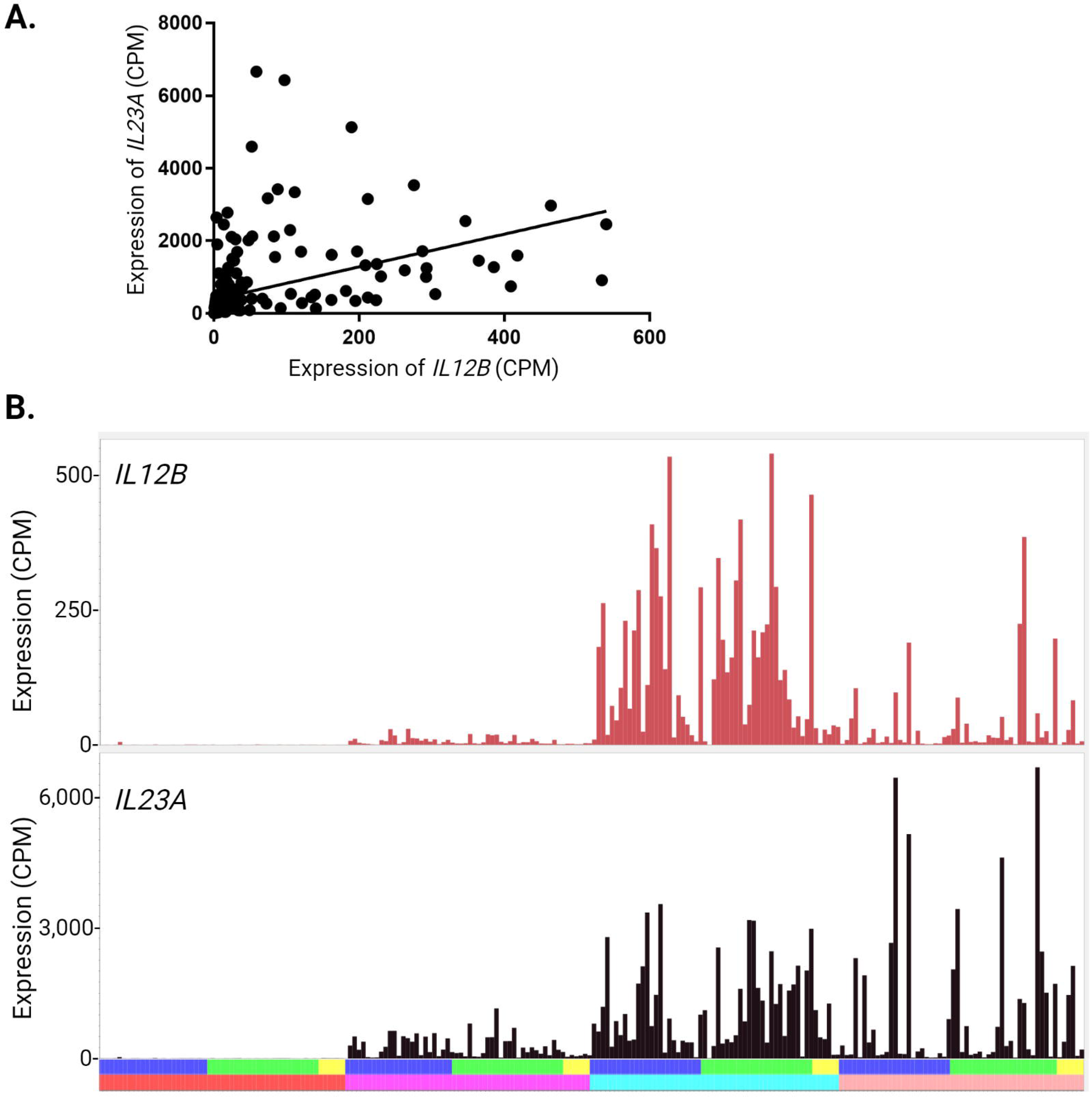
Expression of *IL23A* and *IL12B*. *IL23A* and *IL12B* encode the two subunits of the heterodimeric cytokine, IL23. Expression of the two genes was extracted from RNA-seq analysis of the time course of the LPS response of MDM from 56 individuals (primary data in **Table S1**) *A. Correlation between IL23A and IL12B expression*. Graph shows the relationship between the two transcripts across the full dataset (all 4 time points) including the line of best fit. Spearman *r* = 0.8766, *p* < 0.0001. Zero time values are clustered at left. Note the numbers of outliers above (high *IL23A*) and below (high *IL12B*) the line. *B. Patterns of IL23A and IL12B expression across the time course*. Y axis – gene expression in CPM. X axis – samples ordered by time point and disease status; each column represents a single sample. Upper bar – disease status (blue: affected (Crohn’s disease or ulcerative colitis); green: unaffected sibling of an affected individual; yellow: healthy donor with no family history of inflammatory bowel disease). Lower bar – time point (red: 0 hrs; pink: 2 hrs; turquoise: 7 hrs; salmon: 21 hrs).

In humans, *CSF1* mRNA is not detectable in freshly-isolated monocytes but is induced during monocyte-macrophage differentiation. MDM become autocrine for this growth factor [18]. *CSF1* was indeed highly-expressed in unstimulated cells in most individuals and further-induced around 3-fold by LPS at 2hrs, but expression varied independently of other cytokines. A relatively common SNV variant at the *CSF1* locus is linked to Paget’s disease [43] but we found no association between SNVs at the *CSF1* locus and the level of *CSF1* mRNA in MDM (not shown). One feature of the LPS response in human MDM that has not previously been reported is the induction of the chemokine receptor, *CCR7* and its ligand, *CCL19.* Both were variable between individuals and not correlated with each other (**Table S1B**).

Transcripts encoding the three subunits of complement component C1q (*C1QA, CIQB, C1QC*) are also undetectable in monocytes and highly-expressed in MDM. C1q has multiple functions in autoimmunity and in resistance to infection [44]. Each *C1Q* transcript varied more than 10-fold between individuals. *C1QA* and *C1QC* were correlated with each other (Cluster 443) at r>0.75. *C1QB* was correlated at r=0.41 (with *C1QC*) and r=0.54 (with *C1QA*). Potter *et al.* [45] provided evidence of increased synthesis and catabolism of C1q in IBD patients. An IBD susceptibility locus, defined by rs12568930, is located at Chromosome 1:22,375,738. The nearest gene is *ZTBT40,* but the three C1Q genes lie within the genomic interval [18]. Our data suggest C1q provides stronger candidates than others in the interval.

### eQTL analysis

The large majority of genes expressed in human monocytes show evidence of heritable variation in expression, either in the basal state or following stimulation with LPS or interferon-gamma [22, 35, 46]. The relatively small population analyzed herein, and the obvious and deliberate relatedness, is not optimal for any genome-wide association analysis, including eQTL [47]. However, at least some of the variation in gene expression we detected suggested the presence of null expression alleles that might be linked to common SNVs. To test this possibility, we genotyped each of the individuals using a focused immune chip and performed eQTL analysis. **Figure 4** shows the Manhattan plots for each of the 4 time points and **Table S3** contains all associations with a p value of <10^-6^. A major peak of *cis*-acting variation was detected over the HLA region of chromosome 6 at every time point, but otherwise the weaker candidate eQTL associations in **Table S3** are predominantly *trans*-acting.

**Figure 4.**
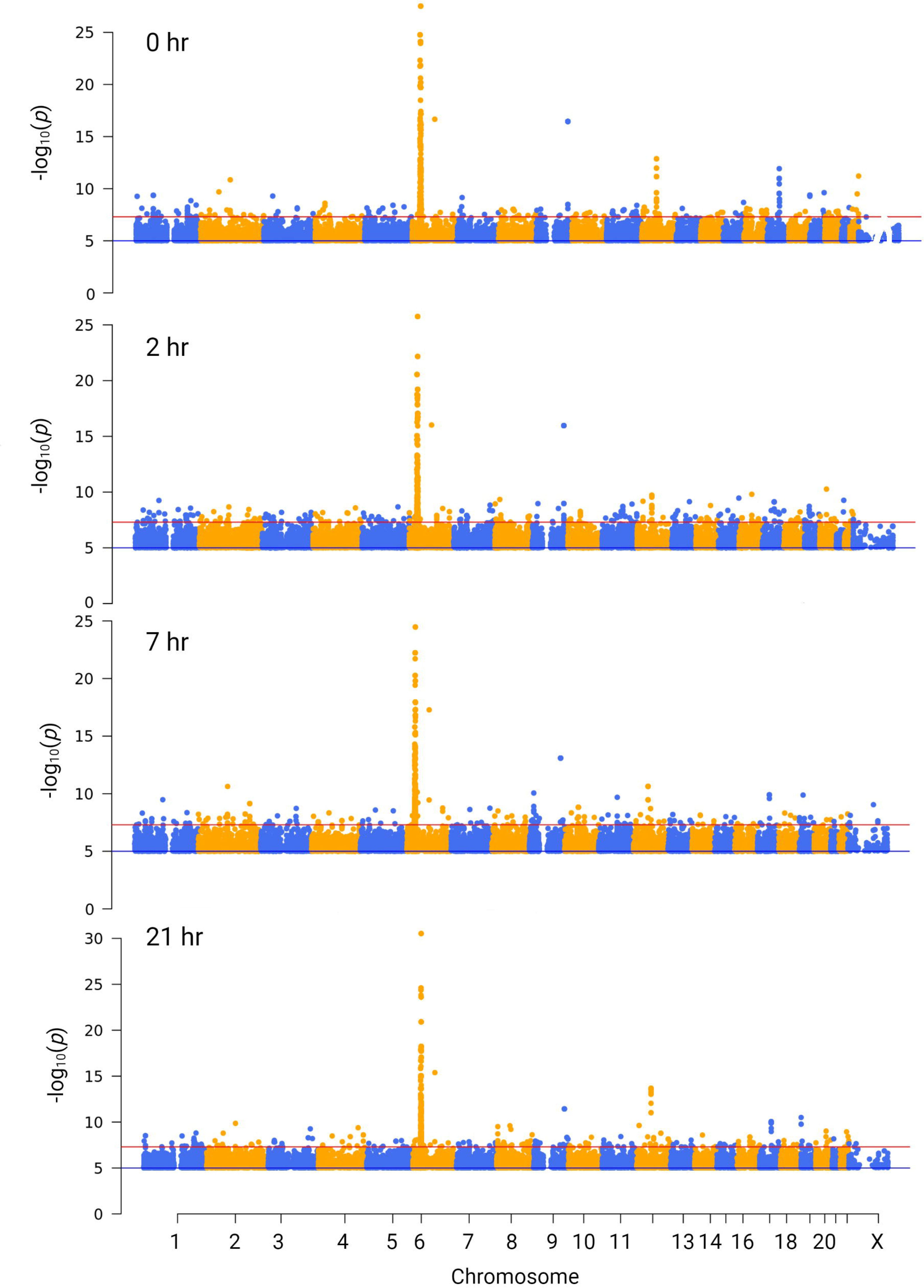
Expression QTL analysis of the response to MDM to LPS. Gene expression profiles from RNA-seq analysis of the time course of the LPS response of MDM from 56 individuals (primary data in **Table S1**) were correlated with SNV genotype as described in Materials and Methods. Y axis – p value (-log10) of the correlation co-efficient for the SNV genotype. Red line shows the p value considered significant after correction for multiple testing. X axis – chromosome number and position for each analysed SNV. Colours alternate with odd numbered chromosomes in blue and even numbered chromosomes in yellow.

### Association of the regulation of MHC gene expression with HLA haplotypes

Because of the extensive polymorphism of HLA and known variation associated with specific haplotypes, it is possible to determine the HLA genotype based upon expressed SNVs [48] and in turn to assess the relationship between HLA type and expression of specific HLA transcripts. We also used the published analytical pipeline RNA2HLA to establish the HLA types of each of the individuals in this cohort to confirm family relationships and sample identity and to link variation to specific haplotypes. In the cluster analysis, the class II MHC regulator *CIITA*, and chaperone, *CD74*, were not correlated with class II MHC gene expression. *CD74* was highly and relatively uniformly expressed and clustered only with *HLA-DPA1 (***Cluster 870**). The only co-regulated HLA genes were pairs located adjacent in the genome, *HLA-B/HLA-C, HLA-A/HLA-F* and *HLA-DRB1/HLA-DRB5*, correlated with each other because of extreme variation between individuals. The expression of these gene pairs across the whole data set is shown **Figure 5**, sorted by family and separately by disease status. In each case, there are siblings with >100-fold differences in expression, with examples of concordant and discordant sib-pairs that are not related to disease status. However, in overview, all but 18 of the 56 tested individuals over-expressed at least one of these HLA gene pairs in MDM. **Table 1** relates the HLA genotypes of each individual to the level of expression of four of these HLA genes. Expression of each of the correlated pairs was not strictly-related to HLA genotype across the cohort, but was correlated with HLA genotype in sib-pairs.

**Figure 5.**
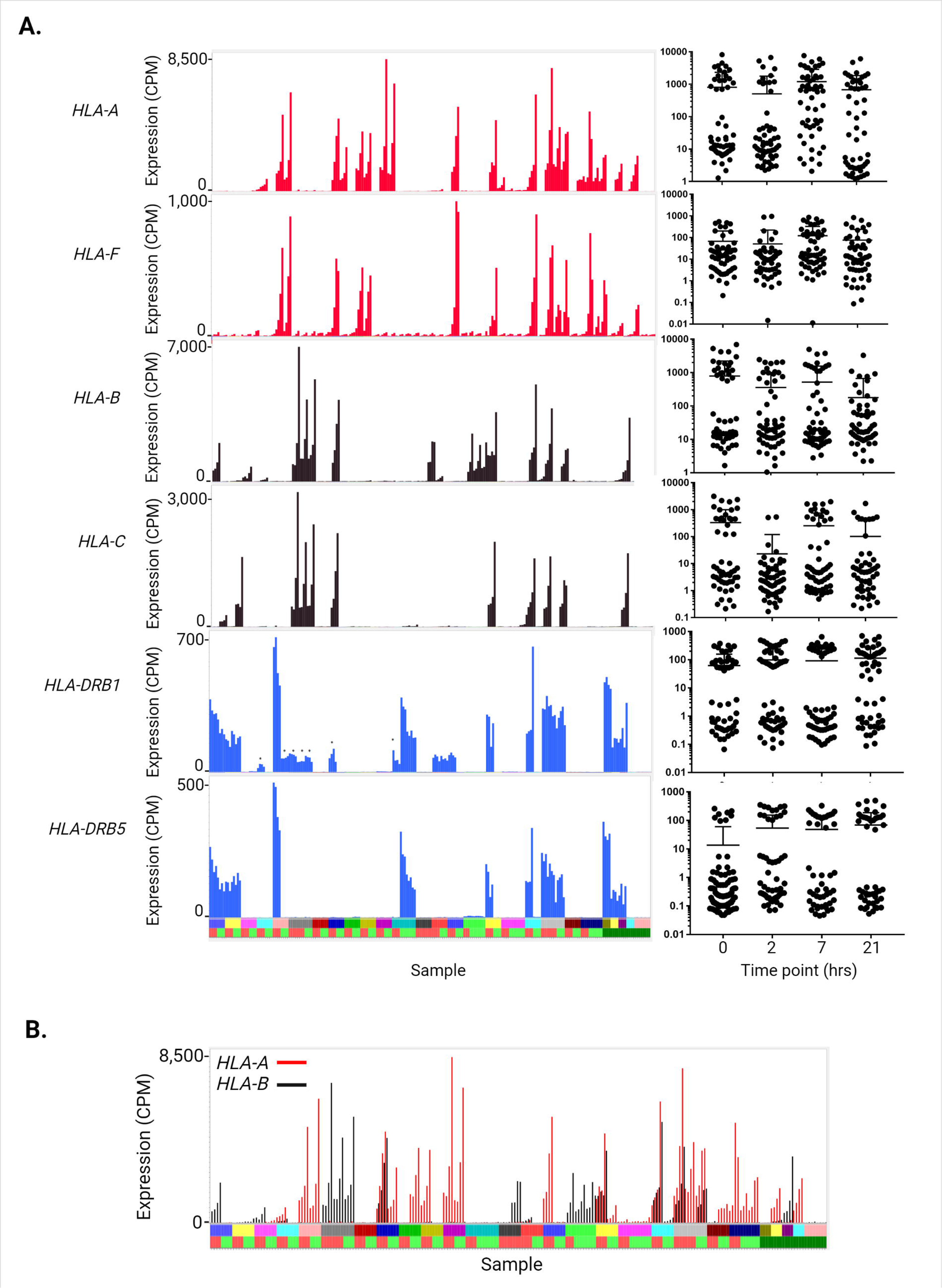
Expression of MHC genes in IBD patients and families. Expression of the Class I and Class II MHC genes was extracted from RNA-seq analysis of the time course of the LPS response of MDM from 56 individuals (primary data in **Table S1**) *A. Expression of MHC genes*. Histograms on the left show the levels of expression for each individual across the four time points. Y axis – expression in CPM. X axis – individuals and time points. Each column represents one sample and the four time points for each individual are adjacent to each other and next to other family members. Expression of *HLA-A* and *HLA-F* (red), *HLA-B* and *HLA-C* (black) and *HLA-DRB1* and *HLA-DRB5* (blue). Panels on the right show the distribution of expression values for the four time points. Y axis – expression (TPM, log_10_ scale). X-axis – time point. Upper bar – family (arranged as in **Table S1**). Lower bar – disease status (red: affected with Crohn’s disease or ulcerative colitis; green: unaffected sibling; dark green – healthy donor with no family history of IBD. *B. Coexpression of Class I MHC genes*. Histogram shows expression of *HLA-A* (red) and *HLA-B* (black) in each individual. Y axis – expression in CPM. X axis – individuals and time points. Each column represents one sample and the four time points for each individual are adjacent to each other and next to other family members. Upper bar – family (arranged as in **Table S1**). Lower bar – disease status (red: affected with Crohn’s disease or ulcerative colitis; green: unaffected sibling; dark green – healthy donor with no family history of IBD. Some individuals express one or other pair of genes, others express both and some express neither. In several families, members show different patterns (boxes).

**Table 1:** HLA genotypes and level of expression of HLA genes. HLA genotypes derived from RNA2HLA (see Methods). Expression levels are given as indicated for each gene. For HLA-A, <250 CPM – 0, 250 CPM≤x<1000 CPM – +, 1000 CPM≤x<3000 CPM – ++, ≥3000 CPM – +++. The maximum value was 8227 CPM and the time point for maximum expression was 21 hrs. For HLA-B, <250 CPM – 0, 250 CPM≤x<1000 CPM – +, 1000 CPM≤x<3000 CPM – ++, ≥3000 CPM – +++. The maximum value was 6948 CPM and the time point for maximum expression was 21 hrs. For HLA-C, <100 CPM – 0, 100 CPM≤x<500 CPM – +, 500 CPM≤x<1500 CPM – ++, ≥1500 CPM – +++. The maximum value was 3132 CPM and the time point for maximum expression was 21 hrs. For HLA-DRB1, <25 CPM – 0, 25 CPM≤x<100 CPM – +, 100 CPM≤x<300 CPM – ++, ≥300 CPM – +++. The maximum value was 704 CPM and the time point for maximum expression was 0 hrs for most individuals.

| Individual ID | Status | HLA-A allele1 | HLA-A allele2 | Max HLA-A expression | HLA-B allele1 | HLA-B allele2 | Max HLA-B expression | HLA-C allele1 | HLA-C allele2 | Max HLA-C expression | HLA-DRB1 allele1 | HLA-DRB1 allele2 | Max DRB1 Expression |
| --- | --- | --- | --- | --- | --- | --- | --- | --- | --- | --- | --- | --- | --- |
| MN1AF | AF | A*01:01 | A*02:01 | 0 | B*08:01 | B*40:01 | ++ | C*03:04 | C*07:01 | 0 | DRB1*15:01 | DRB1*13:02 | +++ |
| MN1UN | UN | A*01:01 | A*68:01 | 0 | B*08:01 | B*44:02 | 0 | C*07:04 | C*07:04 | + | DRB1*15:01 | DRB1*11:01 | ++ |
| MN5AF | AF | A*26:01 | A*02:01 | 0 | B*13:02 | B*52:01 | 0 | C*06:02 | C*12:02 | 0 | DRB1*15:02 | DRB1*07:01 | ++ |
| MN5UN | UN | A*02:03 | A*01:01 | 0 | B*13:02 | B*38:02 | 0 | C*06:02 | C*07:02 | +++ | DRB1*16:02 | DRB1*07:01 | ++ |
| MN6AF | AF | A*02:05 | A*02:05 | 0 | B*50:01 | B*15:01 | + | C*18:01 | C*06:02 | 0 | DRB1*13:01 | DRB1*07:01 | 0 |
| MN6UN | UN | A*24:02 | A*02:01 | 0 | B*15:01 | B*15:01 | 0 | C*04:01 | C*03:03 | 0 | DRB1*13:01 | DRB1*03:17 | 0 |
| MN8AF | AF | A*25:01 | A*11:01 | + | B*39:01 | B*35:01 | + | C*04:01 | C*12:03 | 0 | DRB1*01:03 | DRB1*07:01 | + |
| MN8UN | UN | A*29:02 | A*02:01 | 0 | B*44:02 | B*44:02 | 0 | C*05:01 | C*16:01 | 0 | DRB1*04:01 | DRB1*07:01 | 0 |
| MN9AF | AF | A*03:01 | A*29:02 | +++ | B*44:03 | B*15:01 | 0 | C*03:04 | C*16:01 | 0 | DRB1*15:01 | DRB1*07:01 | +++ |
| MN9UN | UN | A*03:01 | A*29:02 | +++ | B*44:02 | B*40:01 | 0 | C*05:01 | C*16:01 | 0 | DRB1*01:03 | DRB1*07:01 | + |
| MN11AFC | AF | A*31:01 | A*02:01 | 0 | B*39:01 | B*07:02 | +++ | C*12:03 | C*07:02 | +++ | DRB1*01:03 | DRB1*04:04 | + |
| MN11AFU | AF | A*31:01 | A*02:01 | 0 | B*39:01 | B*07:02 | +++ | C*12:03 | C*07:02 | +++ | DRB1*01:03 | DRB1*04:04 | + |
| MN11UN | UN | A*31:01 | A*02:01 | 0 | B*39:01 | B*07:02 | +++ | C*12:03 | C*07:02 | +++ | DRB1*01:03 | DRB1*04:04 | + |
| MN12AF | AF | A*01:01 | A*01:01 | 0 | B*08:01 | B*08:01 | 0 | C*07:01 | C*07:01 | 0 | DRB1*03:01 | DRB1*03:01 | 0 |
| MN12UN | UN | A*01:01 | A*01:01 | 0 | B*08:01 | B*08:01 | 0 | C*07:01 | C*07:01 | 0 | DRB1*03:01 | DRB1*03:01 | 0 |
| MN13AF | AF | A*03:01 | A*02:01 | +++ | B*44:02 | B*07:02 | +++ | C*07:02 | C*05:01 | +++ | DRB1*01:03 | DRB1*04:01 | ++ |
| MN13UN | UN | A*11:01 | A*11:01 | ++ | B*44:02 | B*35:01 | 0 | C*04:01 | C*05:01 | 0 | DRB1*04:08 | DRB1*14:103 | 0 |
| MN14AF | AF | A*01:01 | A*01:01 | 0 | B*08:01 | B*08:01 | 0 | C*07:01 | C*07:01 | 0 | DRB1*03:01 | DRB1*03:01 | 0 |
| MN14UN | UN | A*01:01 | A*03:01 | +++ | B*08:01 | B*35:03 | 0 | C*07:01 | C*04:01 | 0 | DRB1*03:01 | DRB1*10:01 | 0 |
| MN15AF | AF | A*03:01 | A*02:01 | +++ | B*18:01 | B*57:01 | 0 | C*06:02 | C*07:01 | 0 | DRB1*11:04 | DRB1*07:01 | 0 |
| MN15UN | UN | A*32:01 | A*02:01 | 0 | B*44:02 | B*15:01 | 0 | C*05:01 | C*03:03 | 0 | DRB1*12:01 | DRB1*04:01 | 0 |
| MN16AF | AF | A*30:02 | A*03:01 | +++ | B*18:01 | B*53:01 | 0 | C*04:01 | C*05:01 | 0 | DRB1*11:01 | DRB1*03:17 | 0 |
| MN16UN | UN | A*30:02 | A*03:01 | +++ | B*18:01 | B*53:01 | 0 | C*04:01 | C*05:01 | 0 | DRB1*11:01 | DRB1*03:17 | 0 |
| MN17AF | AF | A*02:01 | A*03:01 | 0 | B*44:02 | B*15:01 | 0 | C*05:01 | C*03:03 | 0 | DRB1*01:03 | DRB1*04:01 | ++ |
| MN17UN1 | UN | A*01:01 | A*01:01 | 0 | B*08:01 | B*52:01 | 0 | C*07:01 | C*12:02 | 0 | DRB1*03:01 | DRB1*15:02 | +++ |
| MN17UN2 | UN | A*01:01 | A*01:01 | 0 | B*08:01 | B*52:01 | 0 | C*07:01 | C*12:02 | 0 | DRB1*03:01 | DRB1*15:02 | +++ |
| MN18AFC | AF | A*02:01 | A*01:01 | 0 | B*57:01 | B*35:01 | 0 | C*06:02 | C*18:01 | 0 | DRB1*11:01 | DRB1*07:01 | 0 |
| MN18AFU | AF | A*02:01 | A*31:01 | 0 | B*57:01 | B*40:01 | ++ | C*06:02 | C*03:04 | 0 | DRB1*11:01 | DRB1*07:01 | 0 |
| MN19AF | AF | A*24:02 | A*02:01 | 0 | B*49:01 | B*15:01 | + | C*07:01 | C*03:04 | 0 | DRB1*11:02 | DRB1*01:01 | + |
| MN19UN | UN | A*25:01 | A*02:01 | 0 | B*18:01 | B*15:01 | 0 | C*12:03 | C*03:04 | 0 | DRB1*07:01 | DRB1*01:01 | + |
| MN20AF | AF | A*30:04 | A*03:01 | +++ | B*51:08 | B*35:01 | 0 | C*04:01 | C*16:02 | 0 | DRB1*01:01 | DRB1*13:02 | + |
| MN20UN | UN | A*31:01 | A*02:01 | 0 | B*51:01 | B*44:02 | 0 | C*15:02 | C*16:01 | 0 | DRB1*04:07 | DRB1*07:01 | 0 |
| MN21AF | AF | A*68:01 | A*02:01 | 0 | B*51:01 | B*40:01 | ++ | C*16:02 | C*03:04 | 0 | DRB1*08:01 | DRB1*04:07 | 0 |
| MN21UNa | UN | A*68:01 | A*02:01 | 0 | B*51:01 | B*40:01 | ++ | C*16:02 | C*03:04 | 0 | DRB1*08:01 | DRB1*04:07 | 0 |
| MN21UNb | UN | A*68:01 | A*02:01 | 0 | B*51:01 | B*40:01 | ++ | C*16:02 | C*03:04 | 0 | DRB1*08:01 | DRB1*04:07 | 0 |
| MN22AF | AF | A*01:01 | A*03:01 | +++ | B*37:01 | B*07:02 | +++ | C*07:02 | C*06:02 | +++ | DRB1*15:01 | DRB1*14:01 | ++ |
| MN22UN | UN | A*01:01 | A*11:01 | + | B*08:01 | B*35:01 | 0 | C*07:01 | C*04:01 | 0 | DRB1*03:01 | DRB1*14:01 | 0 |
| MN23AF | AF | A*24:02 | A*02:01 | 0 | B*44:02 | B*18:01 | 0 | C*16:04 | C*07:01 | 0 | DRB1*11:04 | DRB1*13:01 | 0 |
| MN23UNa | UN | A*24:02 | A*02:01 | 0 | B*44:02 | B*18:01 | 0 | C*16:04 | C*07:01 | 0 | DRB1*11:04 | DRB1*13:01 | 0 |
| MN23UNb | UN | A*24:02 | A*24:02 | 0 | B*44:02 | B*51:01 | 0 | C*16:04 | C*01:02 | 0 | DRB1*11:04 | DRB1*03:01 | 0 |
| MN24AF | AF | A*24:02 | A*03:01 | +++ | B*57:01 | B*07:02 | +++ | C*06:02 | C*07:02 | +++ | DRB1*15:01 | DRB1*07:01 | +++ |
| MN24UN | UN | A*32:01 | A*02:01 | 0 | B*38:01 | B*40:02 | + | C*02:02 | C*12:03 | 0 | DRB1*10:01 | DRB1*07:01 | 0 |
| MN25AF1 | AF | A*03:01 | A*03:01 | +++ | B*07:02 | B*47:01 | +++ | C*07:02 | C*06:02 | +++ | DRB1*15:01 | DRB1*04:05 | +++ |
| MN25AF2 | AF | A*11:01 | A*03:01 | +++ | B*18:01 | B*47:01 | 0 | C*07:01 | C*06:02 | 0 | DRB1*15:01 | DRB1*04:05 | +++ |
| MN25UN | UN | A*03:01 | A*03:01 | +++ | B*07:02 | B*35:01 | ++ | C*07:02 | C*04:01 | ++ | DRB1*15:01 | DRB1*01:01 | +++ |
| MN26AF | AF | A*01:01 | A*02:01 | 0 | B*44:02 | B*57:01 | 0 | C*06:02 | C*05:01 | 0 | DRB1*04:01 | DRB1*07:01 | 0 |
| MN26UN | UN | A*30:02 | A*31:01 | + | B*35:01 | B*18:01 | 0 | C*04:01 | C*05:01 | 0 | DRB1*04:07 | DRB1*03:01 | 0 |
| MN27AF | AF | A*03:01 | A*24:02 | +++ | B*08:01 | B*47:01 | 0 | C*06:02 | C*07:06 | 0 | DRB1*03:01 | DRB1*03:01 | 0 |
| MN27UN1 | UN | A*03:01 | A*24:02 | ++ | B*08:01 | B*47:01 | 0 | C*06:02 | C*07:06 | 0 | DRB1*03:01 | DRB1*03:01 | 0 |
| MN27UN2 | UN | A*03:01 | A*24:02 | ++ | B*08:01 | B*47:01 | 0 | C*06:02 | C*07:01 | 0 | DRB1*03:01 | DRB1*03:01 | 0 |
| MN2HD | HD | A*01:02 | A*02:01 | 0 | B*08:01 | B*44:02 | 0 | C*05:01 | C*07:01 | 0 | DRB1*15:01 | DRB1*03:01 | +++ |
| MN5HD | HD | A*03:01 | A*24:02 | ++ | B*38:01 | B*35:03 | 0 | C*04:01 | C*12:03 | 0 | DRB1*15:02 | DRB1*13:01 | ++ |
| MN7HD | HD | A*68:01 | A*24:02 | 0 | B*51:01 | B*07:02 | +++ | C*14:02 | C*07:02 | +++ | DRB1*15:01 | DRB1*10:01 | +++ |
| MN8HD | HD | A*03:01 | A*02:01 | ++ | B*08:01 | B*44:02 | 0 | C*05:01 | C*07:01 | 0 | DRB1*04:01 | DRB1*07:01 | 0 |
| MN9HDa | HD | A*32:01 | A*02:05 | 0 | B*35:03 | B*57:01 | 0 | C*18:02 | C*06:02 | 0 | DRB1*11:01 | DRB1*07:01 | 0 |
| MN9HDb | HD | A*32:01 | A*02:01 | 0 | B*35:03 | B*08:01 | 0 | C*04:01 | C*07:01 | 0 | DRB1*11:01 | DRB1*03:01 | 0 |

### Association of expression with genotype at selected candidate gene loci

Aside from the HLA region, the genome-scale analysis of eQTL does not highlight any clear *cis* SNV associations with the highly-variable expression of individual cytokine genes in **Table S1**. To test the relationship between genotype and expression we examined SNV associations at selected candidate gene loci. *ERAP2* encodes an aminopeptidase involved in antigen processing and lies within a genomic interval associated with IBD susceptibility [18]. There are two major haplotypes with almost equal frequency in human populations [13] one of which encodes a splice isoform that introduces premature stop codons leading to nonsense-mediated decay (NMD). Klunk *et al.* [13] provided evidence that the functional allele was favoured by pathogen selection in survivors of the black death pandemic, caused by the bacterium *Yersinia pestis*, and suggested a consequential increase in susceptibility to inflammatory disease including IBD. **Figure 6A** shows that a subset (13/56) of our cohort has very low expression *ERAP2* in MDM, approximately the proportion of homozygotes anticipated from the allele frequency. In those individuals that expressed *ERAP2* mRNA, expression was not highly variable and was marginally induced by 21hrs. There was no association with disease status within the IBD families. Analysis of expressed SNVs revealed that these individuals were indeed homozygous for the rs1056893 variant (**Figure 6B**). The neighbouring *ERAP1* gene was more highly-expressed than *ERAP2* in MDM and also relatively consistent amongst individuals (**Table S1B-F**). ERAP1 is also involved in peptide processing for presentation on MHC1, and extensive protein-coding variation has been associated with susceptibility to several inflammatory diseases [49]. The calpastatin (*CAST*) locus overlaps *ERAP1* on the opposite strand and has also been implicated in regulation of inflammation in colitis models [50]. *CAST* is highly-expressed by MDM in our data and further induced by LPS. In summary, *ERAP2* is not the only candidate gene within the genomic interval associated with IBD susceptibility.

**Figure 6.**
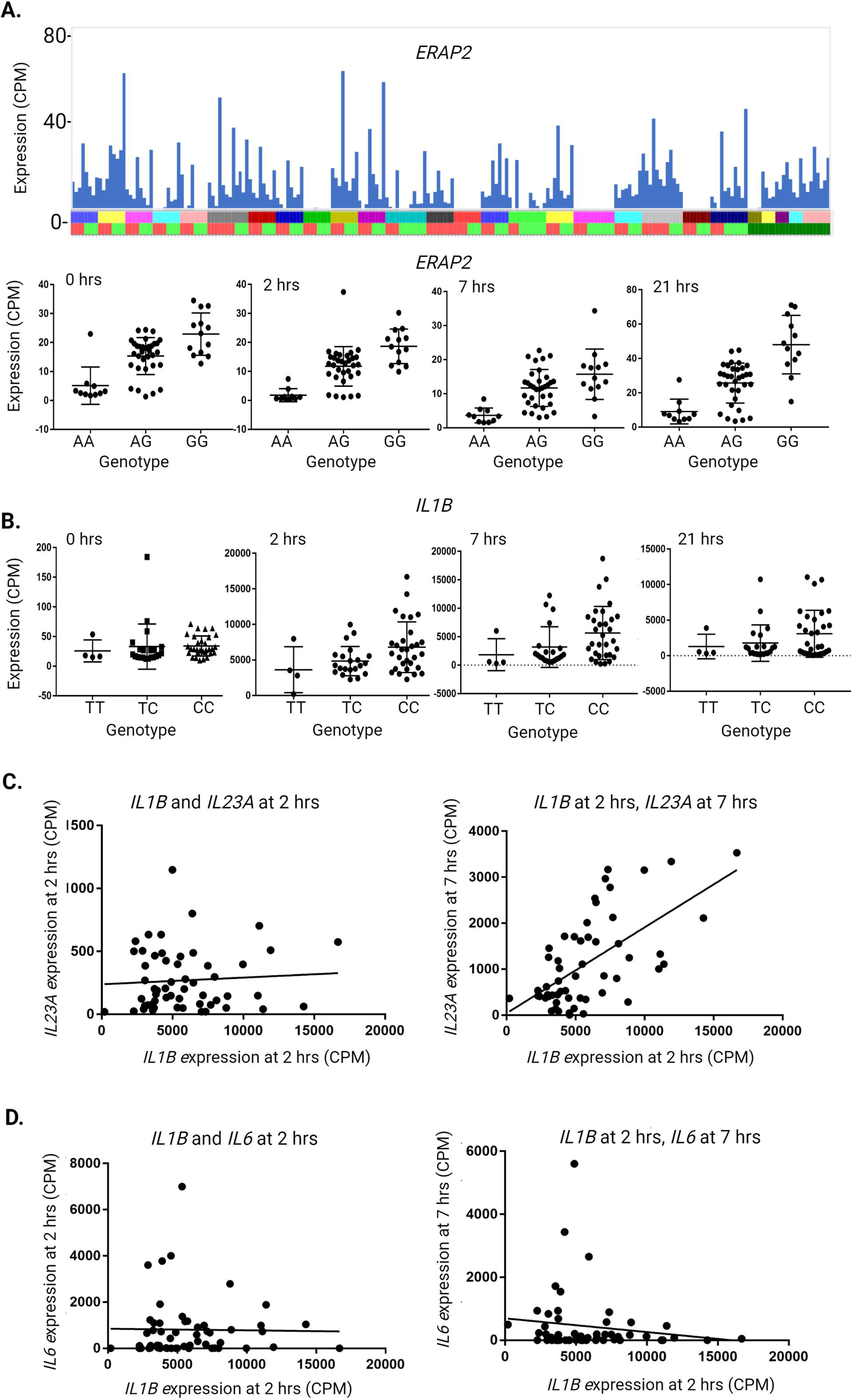
Relationship between SNV genotype and expression of key genes. Expression of the genes indicated was extracted from RNA-seq analysis of the time course of the LPS response of MDM from 56 individuals (primary data in **Table S1**) and SNV genotype was determined as described in Materials and Methods. *A. Expression of ERAP2*. Histogram in the upper panel shows the levels of expression of *ERAP2* for each individual across the four time points. Y axis – expression in CPM. X axis – individuals and time points. Each column represents one sample and the four time points for each individual are adjacent to each other and to other family members. Y axis – expression (CPM). X-axis – genotype. Upper bar – family (arranged as in **Table S1**). Lower bar – disease status (red: affected with Crohn’s disease or ulcerative colitis; green: unaffected sibling; dark green – healthy donor with no family history of IBD). Graphs in the lower panel show the expression values for each genotype at SNV rs1056893 for the four time points. Kruskal-Wallis test showed a highly significant difference among genotypes for all time points (*p* < 0.0001), with the AA genotype showing lower expression than the GG genotype and the heterozygous genotype being intermediate. *B. Expression of IL1B*. Graphs show the expression values for each genotype at SNV rs1143634 for the four time points. Y axis – expression (CPM). X-axis – genotype. Kruskal-Wallis test showed no difference among genotypes for 0 hrs (*p* = 0.17) and 21 hrs (*p* = 39). There was a small difference among genotypes at 2 hrs (*p* = 0.03) and 7 hrs (*p* = 0.02), with the TT genotype showing lower expression than the CC genotype. *C. Correlation between IL1B and IL23A expression*. Left panel shows the relationship between *IL1B* and *IL23A* expression at 2 hrs. Spearman *r* = 0.05577, *p* = 0.68. Right panel shows the relationship between expression of *IL1B* at 2 hrs and *IL23A* at 7 hrs. Spearman *r* = 0.5573, *p* < 0.0001. *D. Correlation between IL1B and IL6 expression*. Left panel shows the relationship between *IL1B* and *IL6* expression at 2 hrs. Spearman *r* = 0.08593, *p* = 0.53. Right panel shows the relationship between expression of *IL1B* at 2 hrs and *IL6* at 7 hrs. Spearman *r* = -0.119, *p* = 0.39.

*IL1B* is of particular interest because of its implication in IBD pathology [51] and because LPS-inducible expression is at least 10-fold lower in MDM compared to freshly-isolated monocytes. Hence, extreme high expression of *IL1B* in response to LPS is an indication of failure of CSF1-induced differentiation. SNV rs1143634 encodes a synonymous substitution with a minor allele frequency (MAF) around 0.2 in most ethnicities (gnomad.broadinstitute.org). That allele frequency is not significantly different in our cohort. **Figure 6B** shows level of expression of *IL1B* mRNA at each time point for each genotype. The distribution of peak expression values is highly skewed; there is a significant correlation between the presence of the minor allele and low expression. Aschenbrenner *et al.* [51] proposed that IL1B provides an autocrine signal regulating expression of another therapeutic target *IL23A* [39, 52] by stimulated monocytes. Consistent with that view, *IL23A* was induced later in the LPS response (peak at 7-21 hrs; **Figure 2**) and expression of *IL23A* at 7 hrs was significantly correlated with *IL1B* at 2 hrs (**Figure 6C**). In contrast there was no correlation between *IL1B* and *IL23A* at 2 hours (**Figure 6C**).

IL6 is another therapeutic target in many inflammatory diseases [53]. *IL6* mRNA at 7hrs varied >100-fold between individuals (**Figure 2**). The *IL6* locus contains an expressed SNV (rs42534390) with an allele frequency of around 50% and a common IL6 promoter variant (-174G/C, rs1800795) has been associated with circulating IL6, variation in responsiveness to LPS and other stimuli [53]. **Figure 6D** shows the relationship between SNV genotype and *IL6* mRNA expression level in our cohort. There was no correlation between *IL6* mRNA and *IL6* SNV genotype, nor with the level of *IL1B* at 2 hrs.

## Discussion

This project was based upon three hypotheses: (1) within human populations there are uncommon, non-coding variants of large effect that influence transcription of key genes in macrophages; (2) differentiation of monocytes in response to CSF1 and a regulated response to microbial stimulus is a key event in adaptation to the pro-inflammatory environment of the lower GI tract; (3) if they exist, regulatory variants affecting basal or inducible gene expression in monocytes differentiated in CSF1 would be prevalent in IBD families. These hypotheses were based upon substantial enrichment identified in large-scale GWAS studies for IBD association with SNVs in the vicinity of the promoters of regulated macrophage-expressed genes [18]. The analysis herein provides further strong support for the view that transcriptional dysregulation in macrophages has a causal relationship to IBD susceptibility. Based upon inducible expression of an NF-κB-regulated luciferase in patient macrophages in a cohort of similar size to ours, Papoutsopoulou *et al*. [54] claimed that CD patients were hyper-responsive to TLR4 signalling. In our data, the overall patterns of gene expression in stimulated macrophages clustered based upon time, reflecting the major changes induced by LPS, but there was no relationship with disease status (**Figure 1B**). Hence, the data provide no support for the idea that IBD arises from a global hyper-responsiveness to TLR4 signals. The available evidence of coding variation in IBD also supports macrophages as the main focus of IBD susceptibility variants. Based upon mass scale genome and exome sequencing in thousands of individuals, Sazonovs *et al.* [11] confirmed known and identified a number of new coding variants apparently enriched in IBD patients. The pattern of expression and function of transcripts containing these coding variants in monocytes and macrophages is summarized in **Table 2.** In overview, the large majority of these coding mutations are likely to impact macrophage function. With the exceptions of *FUT2, IL32R, HGFAC* and *IRGM,* all IBD-associated genes with coding variation identified through genome sequencing are expressed by monocytes and/or MDM and in many cases share the pattern of down-regulation in MDM with *NOD2*, and/or regulation by LPS. In the case of *FUT2* and *IL23R,* variation may impact downstream of macrophage secreted products, IL1B and IL23A. The discussion of roles of *HGFAC* variants by Sazonovs *et al.* [11] focusses on known biology of hepatocyte growth factor (HGF), but *HGFAC* is expressed exclusively in hepatocytes [55] so the mechanistic connection to IBD susceptibility is not obvious. The FANTOM5 analysis implied strongly that *IRGM* in humans is a pseudogene. There is no evidence of expression of *IRGM* mRNA in either the massive CAGE dataset [55] or in the RNA-seq data generated herein. However, active monocyte-macrophage expressed enhancers were detected throughout the *IRGM* locus and flanking DNA. As previously discussed [18] the neighbouring *TNIP1* gene (aka *ABIN3*) encodes a more obvious regulator of inflammation located in this genomic region and regulatory variation in this gene has been detected in patients with systemic lupus erythematosus[56]. *TNIP1* mRNA was highly-expressed in MDM, further induced > 10-fold by LPS and highly variable amongst individuals (**Table S1C-F**).

**Table 2:** Macrophage-related function of candidate IBD susceptibility genes. Genes below were shown to have coding sequence variants significantly linked to IBD susceptibility in large-scale whole genome sequencing analysis of affected individuals and controls [11]. Comments relate to expression in monocytes and macrophages in the current dataset and/or FANTOM5 CAGE data analyzed previously [18]. Selected references highlight known functions of gene of interest and/or neighbouring genes in macrophages or inflammation.

| Gene name | Function/pathway | Comments | References |
| --- | --- | --- | --- |
| *NOD2* | CARD15. Pathogen pattern recognition | Several coding variants. *NOD2* mRNA high in monocytes, down-regulated by CSF1. Co-regulated with *SNX20* and *CYLD* | [87, 88] |
| *ATG16L1* | Autophagy | T300A coding variant. Mechanism/function still unclear. Widely-expressed. Neighbouring *INPP5D* (SHIP1) gene high in monocytes, down-regulated by CSF1. | [89, 90] |
| *SLC39A8* | Zinc transporter (ZIP8) | Monocyte/macrophage specific. Highly-inducible by LPS. Note adjacent *NFKB1*, also inducible by LPS. Highly variable expression | [91, 92] |
| *TYK2* | Tyrosine kinase | Broad expression. Neighbouring gene *ICAM3* high in monocytes, down-regulated by CSF1, implicated in IBD | [93] |
| *IFIH1* | MDA5. Anti-viral defense | Strongly induced by LPS in monocytes and MDM. Variable expression. Link to viral triggers? | [94, 95] |
| *LRRK2* | Multifunctional kinase. PARK8/Dardarin/RIPK7 | High in monocytes and granulocytes. Down-regulated by CSF1. Regulator of phagosome-lysosome homeostasis | [96-98] |
| *SLAMF8* | Cell surface receptor (CD353) | Undetectable in monocytes, induced by CSF1. Inhibitor of reactive oxygen production & inflammation | [99, 100] |
| *PLCG2* | Phospholipase/eicosanoid regulation | Coding mutants in autoinflammatory diseases. Constitutive in monocytes and MDM. Downstream regulator of CSF1 signals | [101] |
| *CARD9* | Signalling adaptor protein | High in monocytes. Repressed by CSF1 and by LPS. Regulation of *IL1B* and inflammasome activation | [102] |
| *ADCY7* | Adenylate cyclase | Monocyte-macrophage specific promoter. Regulates LPS responsiveness. Coding variants associated with ulcerative colitis | [103, 104] |
| *GPR65* | pH-sensing receptor | High in monocytes and granulocytes, Down-regulated by CSF1. Regulator of inflammation | [105] |
| *SMAD3* | Transcription factor | High in monocytes, down-regulated by CSF1. Regulator of inflammation | [106] |
| *IL10RA* | IL10 receptor | High in monocytes, down-regulated by CSF1 but highly-induced by LPS and variable. Mutation in early-onset IBD | [107] |
| *PTAFR* | Platelet activating factor receptor | High in monocytes, down-regulated by CSF1. | [108] |
| *TAGAP* | RhoGTPase activating protein | High in monocytes, down-regulated by CSF1, induced LPS, highly variable. Shared risk with celiac disease. | [109] |
| *IRGM* | Putative regulator of autophagy. | No detectable mRNA in any tissue<br>Neighbouring *TNIP1* (ABIN3) highly-expressed in MDM, induced further by LPS. Binding partner of TNFAIP3 | [110] |
| *RELA* | NFkB transcription factor | Major transducer of LPS signaling. High in monocytes and MDM, further induced by LPS | [54] |
| *DOK2* | Adaptor protein, regulator of tyrosine kinase signals | High in monocytes, down-regulated by CSF1. Further repressed by LPS | [111] |
| *CCR7* | Chemokine receptor | Induced around 500-fold by LPS by 21 hrs. Very variable between individuals. Major ligand, CCL19, also induced >1000-fold. Generation of tertiary lymphoid structures | [112] |

The top 100 hypervariable transcripts at 2, 7 and 21 hrs (**Table S1D-F**) include numerous chemokines (CCL2, CCL4, CCL5, CCL19, CCL20, CCL3L3, CCL4L2, CCL8, CXCL1, CXCL5, CXCL8, CXCL9, CXCL10, CCL15, CXCL3, CCL22, CXCL11) and cytokines (CSF2, CSF3, EBI3, IFNB1, TNF, TNFSF10, IL1A, IL1B, IL6, IL12B, IL23A, IL27, IL32) many of which are considered targets for therapeutic intervention. This cocktail of inducible macrophage-derived effectors likely contributes to the formation of tertiary lymphoid tissue that is a feature of IBD [57]. In each case, a simple NCBI search on gene name AND polymorphism identified evidence of an association with disease susceptibility and many of these genes lie within genomic intervals associated with IBD in GWAS analysis (e.g. CXCL chemokines on Chr4 with rs2472649 and CCL chemokines on Chr17 with rs3091316). For each of the cytokines in **Figure 2**, we have also performed pairwise comparison of affected and unaffected siblings as well as global comparisons of affected versus unaffected and healthy donors (not shown). Consistent with the lack of detection of cis-acting variants in the eQTL analysis in **Table S3** we found no evidence for any significant association between magnitude or pattern of expression of any gene and disease. In the specific cases of IL1B and IL6, the analysis in **Figure 6** supports the view that the extensive variation is only partly influenced by cis-acting variants. There has been one report of an association between IBD and a common IL6 promoter polymorphism [58] but we found no evidence of SNV association with the peak level of IL6 mRNA. The IL6 locus has at least nine LPS-responsive inducible enhancers [18] so there is considerable complexity in transcriptional regulation and it somewhat unlikely that a single variant is causal. Given the extensive network of feedforward and feedback regulation analysed previously [18] expression of each transcript is essentially a complex trait. Identification of cis or trans-acting regulatory variants of large effect would be even more challenging than the search for coding variants. Functional annotation of non-coding SNVs would be even more challenging. The published analysis of HLA sequences identified thousands of non-coding variants within putative regulatory elements [56].

Aside from cytokines and chemokines, each hypervariable regulated transcript expressed in MDM can form the basis for a hypothetical link to IBD susceptibility. For example, *IFITM1, IFITM2* and *IFITM3* contribute to endocytic targeting of incoming virus particles, restricting virus replication [59]. Each of these transcripts is highly-expressed by monocytes and down-regulated by CSF1, enabling entry of influenza virus, expression of viral proteins and induction of pro-inflammatory genes [60]. *IFITM3* variants are associated with severe disease in influenza infection and an *Ifitm3* knockout mouse is susceptible to low pathogenicity infections [61]. Mice lacking the entire IFITM locus develop spontaneous colitis [62]. In unstimulated MDM, *IFITM1* was barely detectable but there was residual expression of *IFITM2* and *IFITM3.* Each transcript was induced by LPS with differences in time course and magnitude, and large variations between individuals, but they were sufficiently correlated to form part of the interferon-response **Cluster 3** in the current data set. Could the extreme variation in expression of these and other LPS-inducible interferon-responsive genes (*CXCL10, IDO1, IFIT2, ISG15, ACOD1, RSAD2, OSAL, IFIH, ISG20*) predispose to hyper- or hypo-responsiveness to enteric viruses as the initial environmental trigger of chronic inflammation?

The relative abundance and function of inducible negative feedback regulators of macrophage activation highlighted previously [18] also varied between individuals. For example, the set of highly-expressed and hypervariable transcripts in **Table S1D-F** includes *ACOD1, DUSP1, GPR183, OTUD1, PMAIP1, PTGS2 (aka COX2), SERPINE1, SOCS3, TNFAIP3, TNFAIP6* and *ZFP36*. These have been described as inflammation suppressor genes, in that even heterozygous mutations in any one of them can lead to spontaneous inflammation [63, 64]. Variation in their expression is likely to contribute to the variation in temporal profiles of cytokine induction, for example the transient or sustained expression of IL1B (**Figure 2**). In mice, transcripts encoding anti-inflammatory IL10 (*Il10* gene) and IL10 receptor (*Il10ra* gene) appear to be required for homeostatic maintenance and regulation of IL23 production but the available evidence implicates T cells as the main source of IL10. Conditional deletion of either ligand or receptor in macrophages did not lead to spontaneous inflammation but influenced the response to pathogenic bacterial challenge [65]. *IL10* mRNA was induced in MDM by LPS in all individuals but the level of expression was relatively low and both the time course and the peak expression were highly variable (see **Table S1D-F,** **Figure 2**). The receptor gene, *IL10RA,* was expressed in unstimulated MDM and induced 6-7 fold further by 7 hrs following LPS addition (**Table S1E**). This suggests that main axis of control is the regulated response of macrophages to IL10 produced by other cells. SNVs in both *IL10* and *IL10RA* have been linked to IBD susceptibility in GWAS and loss-function mutations are associated with very early onset disease [66].

The profound induction by LPS of transcripts encoding the chemokine receptor, CCR7, and its major ligand, CCL19, has not been reported previously. At each of the time points, *CCR7* expression varied between individuals (**Table S1C-F**). The literature on CCR7 biology focusses exclusively on its expression by dendritic cells (DC) and the function in guiding these cells to draining lymph nodes (e.g. [67]). Transcripts encoding inducible co-stimulatory/inhibitory molecules detected on active human DC [68], CD83, CD80, CD40 and CD274 (PDL-1) were inducible >20 fold by LPS in MDM and highly variable between individuals. The macrophages of the lamina propria of mice, conflated with dendritic cells because of their expression of CD11c (*Itgax*) (see [69]) express CCR7 and migrate in response to CCL21 [70]. As expected, the human monocytes differentiated in CSF1 expressed *ITGAX* (CD11c) and *ITGAM* (CD11b) mRNA. They also express *ITGAE* (CD103), commonly used as a marker of mouse and rat DC [69], but lack detectable FLT3. Studies in the rat have demonstrated that migrating “dendritic cells” in lymphatics draining the gut are derived from blood monocytes [71]. We conclude that monocytes differentiated in CSF1 can be primed to respond to microbial agonists by becoming migratory antigen-presenting cells (APC).

Of course, the primary determinant of APC activity is the expression of MHC molecules to enable presentation of antigenic peptides to T cell receptors. If monocyte differentiation in CSF1 provides a model for adaptation to the gut mucosa [18], then the differential expression of both Class I and Class II MHC genes, independent of their antigen binding specificity, may well underlie the selective response to classes of mucosal antigens leading to inflammation. As noted in the introduction, shared haplotypes in the HLA region provide the greatest prediction of shared risk of IBD amongst members of the same family [1, 72]. High-density SNV typing of the HLA region in >32,000 individuals with IBD implicated several HLA alleles, with the greatest effect size associated with HLA-DRB1*01:03 in both CD and ulcerative colitis [73]. Seven of the individuals in our cohort (five affected, two unaffected) have this allele, all as heterozygotes. This included three members of one family (MN11), with one unaffected individual, one with CD and one with ulcerative colitis. The discussion of these specific HLA associations has focused on variation in the antigen-binding sites, with the presumption that these variants predispose in some way to response to an environmental trigger. There is also evidence of associations of class I MHC (specifically HLA-C) alleles with IBD [74] and of regulatory polymorphism affecting HLA-C expression [75]. High HLA-C expression on peripheral blood leukocytes was associated with increased risk of CD [76]. 11/50 individuals in our cohort expressed high levels of HLA-C (>400TPM) in MDM (compared to an average of <2TPM in non-expressors) but there was no correlation with disease status (**Figure 6**). By contrast to mouse, where class II MHC genes are almost undetectable in macrophages grown in CSF1 (see BioGPS.org), Class II MHC genes are expressed to varying degrees in human blood monocytes and retained or increased in MDM grown in CSF1. The relationships between HLA alleles and expression in relevant cell types has not been explored in the context of IBD. Raj *et al* [56] reported the existence of numerous regulatory polymorphisms affecting expression of Class II MHC genes in monocytes cultured in CSF2 (GM-CSF) (so-called monocyte-derived DC, moDC), as related to susceptibility to systemic lupus erythematosus. However, the effect sizes were small. By contrast, our analysis of MDM grown in CSF1 revealed apparent null expression of class I (HLA-A, HLA-B, HLA-C, HLA-E, HLA-F) and class II (HLA-DRA, HLA-DRB1, HLA-DRB5, HLA-DQA1, HLA-DQB1, HLA-DMB, HLA-DPA1) in certain individuals, linked to HLA haplotypes (**Table 1**). HLA-C is of particular interest as the major determinant of natural killer (NK) cell recognition [77, 78]. The small number of individual monocytes and MDM profiles in the FANTOM5 project also show >10-fold variation in expression of each of these genes among individuals. These large differences could regulate acquired immune responses to broad classes of microflora that might in turn initiate a chronic inflammatory state.

## Conclusions

This study has not identified a common global pattern of dysregulated variation in macrophage activation that distinguishes affected and unaffected individuals or discordant sib-pairs. What it has done is to confirm that in macrophages differentiated in CSF1 there are very large inter-individual differences in basal and LPS-inducible expression of specific pro-inflammatory or anti-inflammatory genes. Whereas analysis of common SNVs by GWAS reveals variants of small effect on disease susceptibility at a population level, we suggest that within individual families, there are smaller numbers of regulatory variants of much larger effect that impact gene expression in macrophages and could only be associated with disease through conventional linkage analysis in large pedigrees. The concept that each family is unique has been termed the Anna Karenina hypothesis for the genetics of complex disease [79]. The corollary of the view is that IBD may, in fact, be many different diseases with unique underlying genetic basis and corresponding variation in treatment efficacy.

## Materials and methods

All patients provided consent and the study was approved by the ACT Health Human Research Ethics Committee (ETH.5.07.464).

### Blood monocyte isolation

Approximately 65 mL of blood was drawn from each subject into ACD-B Vacuette® tubes, transferred to sterile 50 mL tubes, and centrifuged for 30 min at 12,000 x g with no brake. A 200 uL aliquot of blood was kept for DNA extraction. Plasma was removed from the tubes so that only a 1 cm layer remained. Approximately 9 mL of buffy coat was aspirated from each sample and diluted 1/1.5 in RPMI with no additives. The buffy coat dilution was gently layered on to Lymphoprep™ to a total volume of 50ml in a 50mL tube, then spun for 45 min at 200 x g at room temperature with no brake. The clear top layer was removed and the mononuclear cells (peripheral blood mononuclear cells; PBMC) was recovered, diluted in 30ml RPMI1640 and pelleted by centrifugation. Cells were counted, washed once with phosphate buffered saline. CD14-positive cells were enriched by using the Classical Monocyte Isolation Kit (Miltenyi 130-117-337) according to the Manufacturer’ instructions. A 10 uL aliquot of the magnetic column eluate was removed for cell counts and purity check. The remainder of the elution was centrifuged for 10 min at 300 x g at 4°C.

### Cell culture

CD14^+^ monocyte pellet was resuspended in complete culture medium; RPMI1640 (Sigma-Aldrich) containing 2mM Glutamax (Gibco), 20ug/ml penicillin/streptomycin (Sigma-Aldrich), and 10% v/v human AB serum (Sigma-Aldrich). Recombinant human macrophage colony-stimulating factor (CSF1, Gibco-Thermo Fisher, PHC9501) was then added at a final concentration of 100 ng/ml. The cells were then plated at 10^6^ cells per well, to a final volume of 2 mL. The 6-well plates were coated with 1ug/cm^2^ human plasma fibronectin (Sigma-Aldrich) as per the manufacturer’s instructions to create a more physiologically relevant substratum. The 6-well plates were placed in the incubator at 37°C with 5% CO_2_. At day four of incubation 1 mL of fresh complete medium with 3 X CSF1 concentration was added to each well. On day six, the medium was replaced with 2 mL complete medium with 1X concentration of CSF1. On day seven of incubation, the time course experiments commenced (Day 1), LPS 10 ng/ml lipopolysaccharide *Salmonella enterica* serotype Minnesota Re 595 (Sigma-Aldrich) was added to each well. Time point 0 hr was collected before LPS was added. At each time point (2 hr, 7 hr, 21 hr), cells were lysed using 0.5 mL TRIzol (Invitrogen), after which 100 uL chloroform was added. The plates were rocked back and forth for 15 sec and incubated at 37°C with 5% CO_2_ for 2 min. The cells were then placed in sterile microcentrifuge tubes and placed at -80°C.

### Transcriptome sequencing

A total of 224 samples underwent whole genome transcriptome sequencing. Lysed cell samples were thawed and RNA extracted using a Qiagen RNeasy miniprep kit, including DNase digestion (Qiagen). RNA concentration and purity was assessed on a Tapestation (Agilent). Library preparation was performed using Illumina TruSeq Stranded mRNA kits. Sequencing was performed on a NovaSeq 6000 instrument using S1 flowcells (three in total) in a 100 bp paired-end format. Library preparation and sequencing were performed by the Biomolecular Resource Facility of the Australian National University, Canberra, Australia, according to manufacturer’s instructions.

### SNV genotyping

DNA was extracted from a 200 uL blood sample for each subject using QIAamp DNA Blood Mini kits with RNaseA digestion (Qiagen). SNV genotyping was performed using Illumina Infinium ImmunoArray-24 v2.0 BeadChips (InfiniumImmunoArray-24v2-0; Illumina, Inc., San Diego, CA, USA). A total of 4 uL DNA was loaded onto the array, as per the manufacturer’s guidelines. The BeadChips were scanned using an Illumina iScan. Samples were genotyped using Illumina’s GenomeStudio 2.0.4 with Genotyping module 2.0.4 software, using default settings. The Illumina InfiniumImmunoArray-24v2-0 manifest file and project specific cluster file were used for the analysis (InfiniumImmunoArray-24v2-0_A_ClusterFile.egt). SNV genotyping was performed by Australian Genome Research Facility, Melbourne, Australia.

### Bioinformatics

FastQC (https://www.bioinformatics.babraham.ac.uk/projects/fastqc/) was used to assess the quality of RNA-seq results. Assignment of reads to transcripts was then conducted using the pseudoaligner Kallisto v0.44 [80]. Ensembl release 103 of the GRCh38 genome was used for all kmer matching and subsequent annotation. The R package tximeta [81] was used to calculate gene-level expression, corrected for effective gene length, resulting in a file for abundance, in counts per million (CPM). Ensembl gene and primary transcript IDs, NCBI stable gene IDs, Ensembl gene symbols and HGNC symbols were included for each gene. This preparation of the data for analysis was carried out by the ANU Bioinformatics Consultancy, Australian National University, Canberra, Australia.

Network analysis was then performed on the abundance file. Lowly expressed genes (CPM<1.0) and genes expressed in <2 sample records (N records = 222; N samples = 56) were removed. The resultant dataset consisted of 19,972 genes. The network analysis tool BioLayout (http://biolayout.org, formerly BioLayout *Express*^3D^ [30]) was used to visualize the transcriptomic data and explore relationships between samples (sample-to-sample analysis) and genes (gene-to-gene analysis). For the sample-to-sample analysis, a threshold Pearson correlation coefficient of 0.87 was used; this was the maximum value that included all samples. For gene-to-gene analysis, a Pearson correlation coefficient of 0.7 was chosen to maximise the number of genes included while minimizing the computational complexity due to the number of connections between them [82].

Analysis of the SNV genotyping data was performed by Australian Genome Research Facility, Melbourne, Australia. The hg38 build of the human reference genome was used for analysis. Final reports were merged and converted to PLINK PED/MAP format using gcta v1.91.2 (Yang et al., 2011). Alleles were flipped to the positive strand using the ImmunoArray v2.0 strand designations (N=102,173). Quality control was carried out using PLINK [83]. Filtered SNVs were pruned by pairwise genetic correlation to create a set of independent variants appropriate for principle components analysis (PCA). LD pruning was calculated using a window size of 1000 bp and 50 bp step with r^2^ threshold of 0.5. Annotated PCA plots were generated using the ggplot2 package [84] in the R statistical environment. PCA plots were annotated for phenotype, family, age, and sex. Expression QTL (eQTL) analysis was performed for 56 samples using Matrix-eQTL version 2.3 [85] in the R statistical environment version 4.1.2. Four timepoints were modelled in linear mode (least squares), with sex applied as a covariate. A total of 1,194,937,309 eQTL were modelled for each timepoint. A stringent minor allele frequency threshold of 0.2 was applied to reduce inflation generated by rare alleles [86] (approximately 100,000 SNVs remaining in the analysis). To reduce the resultant file sizes, only sites with a significance score of P<1x10^-5^ were exported. Quantile-quantile plots and p-value histograms were generated to assess the effectiveness of the model and data QC.

Following initial QC of SNV results, assignment of the time point samples to individuals was verified and HLA-A, HLA-B, HLA-C and HLA-DRB1 typing was obtained using RNA2HLA [48]. Assignment of sex was validated using expression of Y chromosome genes.

## Data availability

All of the processed RNA-seq data is presented in **Table S1**. The raw RNA-seq sequence files that support the findings of this study are available on reasonable request from the corresponding authors. The data are not publicly available due to privacy and ethical restrictions.

## Acknowledgments

Grant support to PP was provided by Canberra Hospital Private Practice Trust Fund. DAH and KMS are grateful for core support from The Mater Foundation. DAH is supported by NHMRC Investigator Grant 2009750. We thank Dr Zhi-Ping Feng and Mr Cameron Jack of the ANU Bioinformatics Consultancy for initial analysis and quantification of the RNA-seq data. Additional bioinformatic analysis was performed by Dr Lesley Gray at the Australian Genome Research Facility.

## Supplementary Tables

**Table S1. Gene expression data.**

S1A contains the metadata contains metadata describing the individuals and their disease status. Table S1B contains the full set of gene expression based upon quantification using Kallisto as described in Materials and Methods, sorted based upon individual, family and gene name. Tables S1C, S1D, S1E and S1F contain the data for individual time points, 0, 2, 7 and 21 hrs respectively, sorted based upon maximum/minimum expression.

**Table S2. Co-expression clusters**

Table shows the complete set of co-expression clusters generated by network analysis using Biolayout. For each cluster with at least 10 transcripts the Tables shows the average expression profile across the dataset in the same format as in Figure 1B.

**Table S3. Expression QTL analysis**

The Table shows the relationship between SNV genotype and level of expression of specific transcripts at each of the 4 time points for all associations with p value<10^-6^.

